# Routine sub-2.5 Å cryo-EM structure determination of B-family G protein-coupled receptors

**DOI:** 10.1101/2020.08.21.260851

**Authors:** Radostin Danev, Matthew Belousoff, Yi-Lynn Liang, Xin Zhang, Denise Wootten, Patrick M. Sexton

## Abstract

Cryo-electron microscopy (cryo-EM) experienced game-changing hardware and software advances about a decade ago. Since then, there have been gradual and steady improvements in experimental and data analysis methods. Nonetheless, structural analysis of nonsymmetric membrane proteins, such as G protein-coupled receptors (GPCRs), remains challenging. Their relatively low molecular weight and obstruction by the micelle/nanodisc result in marginal signal levels, which combined with the intrinsic flexibility of such complexes creates difficult structural study scenarios. Pushing the performance limits of cryo-EM requires careful optimization of all experimental aspects. To this end, it is necessary to build quantitative knowledge of the effect each parameter has on the outcome. Here, we present in-depth analysis of the influence of the main cryo-EM experimental factors on the performance for GPCR structure determination. We used a tandem experiment approach that combined real-world structural studies with parameter testing. We quantified the effects of using a Volta phase plate, zero-loss energy filtering, objective lens aperture, defocus magnitude, total exposure, and grid type. Through such systematic optimization of the experimental conditions, it has been possible to routinely determine class B1 GPCR structures at resolutions better than 2.5 Å. The improved fidelity of such maps helps to build higher confidence atomic models and will be crucial for the future expansion of cryo-EM into the structure-based drug design domain. The optimization guidelines drafted here are not limited to GPCRs and can be applied directly for the study of other challenging membrane protein targets.

## 1. Introduction

The “revolution” in cryo-electron microscopy (cryo-EM) began a decade ago and provided substantial improvements in performance that were enabled by direct electron detectors and new data processing methods. Since then, there have been gradual technological advances, primarily on the data analysis side (Danev et al., 2019). Nowadays, most of the published single particle reconstructions are in the 2.5-3.5 Å resolution range (Cianfrocco and Kellogg, 2020). Such maps are adequate for unambiguous tracing of the backbone and localization of most sidechains in well-resolved regions, however, depending on the scientific question being asked, the performance may not be sufficient to confidently address the biological problem. For example, understanding how small molecule ligands bind to and modulate receptors often requires the identification of not just direct interactions with the target but also water molecule networks and this is also critical for structure-based drug design. Resolutions better than 2.5 Å, and preferably better than 2.0 Å where the nature of chemical interactions can be determined with precision, are necessary for structure-based approaches to be effective (Anderson, 2003; Armstrong et al., 2006). Ultimately, the level at which regions of a 3D cryo-EM map are resolved will be limited only by the thermal motions of the molecule. To get as close as practically possible to such performance, the experiment must be tuned for maximum signal extraction.

The outcome of a cryo-EM project depends primarily on the biochemical quality and behavior of the sample. Good homogeneity and sufficient concentration are the main factors that determine the success of an experiment (Danev et al., 2019). Therefore, stringent purification and quality control criteria combined with negative stain and cryo-sample screening are essential for achieving good results. Nevertheless, some targets may be difficult to express or natively extract in sufficient quantity or may adopt a preferential orientation in thin ice. Getting a structure from such challenging samples often requires a practical compromise in the cryo-EM approach, such as data collection in thicker ice areas or with a tilted specimen (Tan et al., 2017). This imposes a performance tradeoff that does not leave much room for refinement of the experiment. Here, we present results from well-optimized samples and therefore some of the conclusions may not apply in difficult cases.

In practice, only general guidelines can be proposed for optimal data acquisition parameters. Each research group or cryo-EM facility has adopted a favored set of experimental settings based on their own experience, published results and/or intuitive expectations. Some choices may be limited by the hardware configuration of the microscope, such as the accelerating voltage, the type of detector and the presence of an energy filter, while other settings, such as specimen grid type, defocus range, total exposure and objective lens aperture, are decided by the researcher or the operator. In general, each of these settings may have a small and seemingly negligible effect on the result, but combining several optimal conditions has a cumulative effect and could lead to a noticeable improvement in map resolution and quality. As demonstrated here, and by recent atomic resolution structures (Nakane et al., 2020; Yip et al., 2020), fine-tuning of the experiment is crucial for pushing the performance limits of cryo-EM.

We began our cryo-EM studies of G protein-coupled receptors (GPCR) three years ago by using the Volta phase plate (VPP) (Liang et al., 2017). Active GPCR-transducer complexes are relatively small (~150 kDa) asymmetric assemblies, which at the time were considered to be very challenging for cryo-EM. Data acquisition and image processing methods were not as advanced as they are today, and the capabilities of the conventional defocus phase contrast approach were not fully explored. To increase the probability of success, we decided to employ the VPP and it produced remarkable results on the first try. Thereafter, we continued using the VPP in GPCR projects with great success (Draper-Joyce et al., 2018; Liang et al., 2018a, 2018b). The results were of similar or better quality than results from the conventional approach, which created the subjective impression that the VPP is beneficial for the study of GPCRs (Gao et al., 2019; García-Nafría et al., 2018a, 2018b; Kang et al., 2018; Kato et al., 2019; Koehl et al., 2019; Krishna Kumar et al., 2019; Maeda et al., 2019; Zhang et al., 2017). Meanwhile, with increasing data acquisition throughput and new image processing methods, the quality of results from the conventional defocus approach continued to improve, even for molecules that are smaller than GPCRs (Herzik et al., 2019). Therefore, we decided to empirically reassess the potential benefit of the VPP for the study of GPCRs and conducted a quantitative comparison between the methods using the same sample. To minimize the influence of other factors, datasets with and without the phase plate were collected from the same grid in a single cryo-EM session. All other experimental parameters were kept the same and the VPP/non-VPP datasets could be easily joined to get a reconstruction from the whole dataset for the structural investigation of the complex (Liang et al., 2020). Modification of a single experimental parameter during data acquisition proved to be a very efficient way for conducting structural and methodological studies in parallel. It does not require additional microscope time and the results are from a real-world sample. We used this tandem experiment approach to also evaluate the effect of zero-loss energy filtering and the objective lens aperture. Furthermore, we quantified the effect of defocus amount and total electron exposure by splitting one of the datasets into corresponding subsets.

The results from the quantitative evaluation of the effect of the Volta phase plate, zero-loss energy filtering, objective lens aperture, defocus amount and total exposure are presented below. In addition, we describe our sample optimization efforts that led to a substantial improvement in result quality.

## 2. Results and discussion

The three datasets used in this study are shown in Figure 1. The samples were active state class B1 GPCR complexes PACAP38:PAC1R:Gs (PAC1R) (Liang et al., 2020), Taspoglutide:GLP-1R:Gs (GLP-1R-TAS) (in preparation) and GLP-1:GLP-1R:Gs (GLP-1R-GLP-1) (Zhang et al., 2020). Reconstructions from the complete set of micrographs in each dataset reached global resolutions of 2.7 Å, 2.5 Å and 2.1 Å, respectively (Figures 1A, D and G). Each dataset comprised two subsets of micrographs collected with a modification of a single experimental parameter (Figure 1 middle and right columns, Table 1).

**Figure 1.**
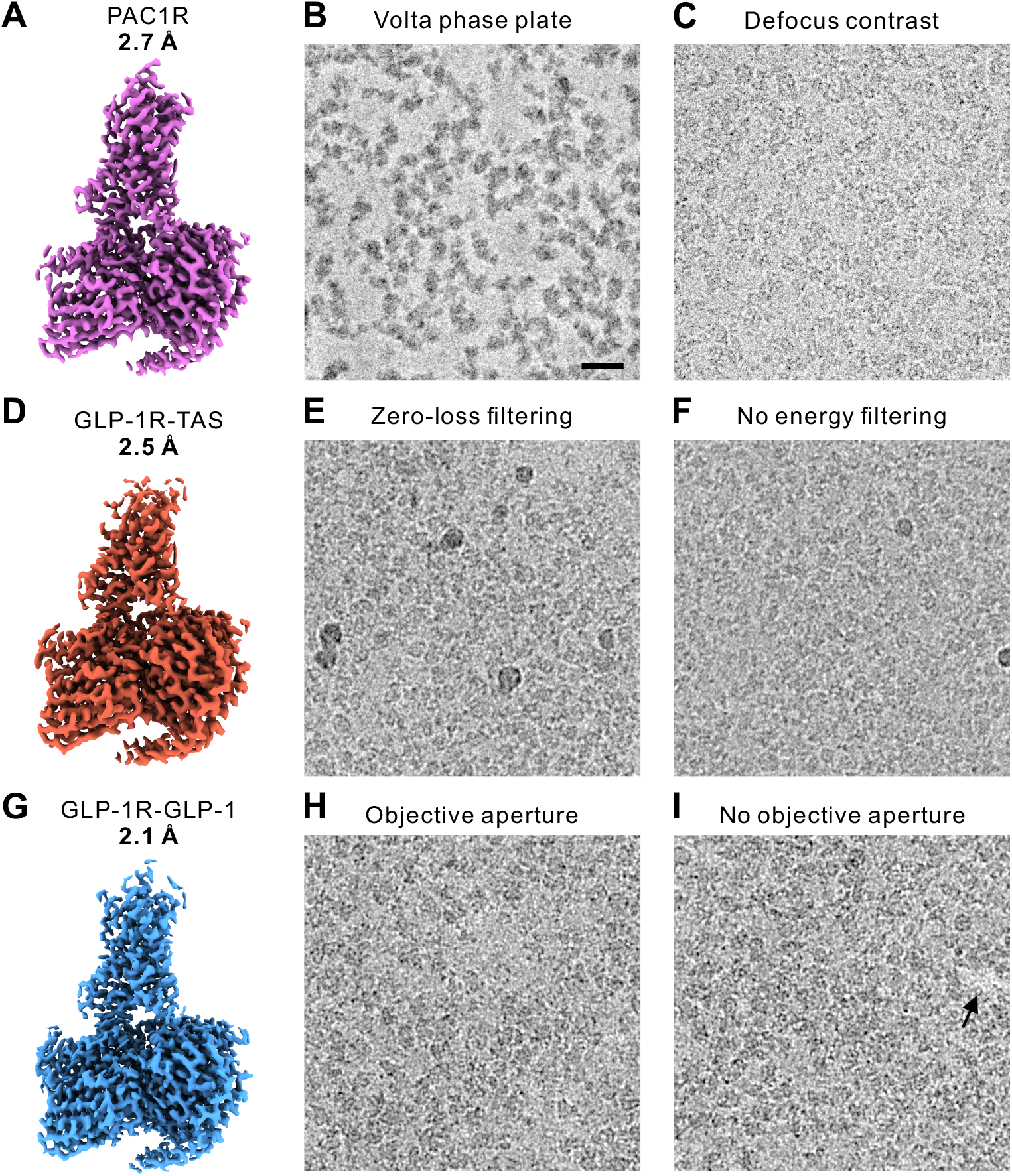
3D maps and representative micrographs from class B GPCR datasets used for the evaluation of experimental parameters. (A) PACAP38:PAC1R:Gs (PAC1R) dataset collected partially with the Volta phase plate (B), and partially with defocus phase contrast (C). (D) Taspoglutide:GLP-1R:Gs (GLP-1R-TAS) dataset acquired partially with zero-loss energy filtering (E), and partially without energy filtering (F). (G) GLP-1:GLP-1R:Gs (GLP-1R-GLP-1) dataset acquired in part with a 100 μm objective lens aperture (H), and in part without an aperture (I). Scale bar 20 nm.

**Table 1.** Cryo-EM experiment details

|  | *PAC1R* |  | *GLP-1R-TAS* |  | *GLP-1R-GLP-1* |  |
| --- | --- | --- | --- | --- | --- | --- |
|  | +VPP | -VPP | +ZLF | -ZLF | +OLA | -OLA |
| **Sample preparation** |  |  |  |  |  |  |
| Concentration [mg/ml] | 5.1 |  | 4.6 |  | 3.7 |  |
| Sample volume [μl] | 3 | | 3 | | 3 | |
| Grid type | Quantifoil<br>R1.2/1.3<br>Cu/Rh 200 |  | Quantifoil<br>R1.2/1.3<br>Cu/Rh 200 |  | UltrAuFoil<br>R1.2/1.3 Au<br>300 |  |
| Glow discharge time [s] | 30 |  | 90 |  | 90 |  |
| Glow discharge current [mA] | 10 |  | 10 |  | 10 |  |
| Blotting chamber temperature [$^{\circ}$C] | 4 | | 4 | | 4 | |
| Blotting chamber humidity [%] | 100 |  | 100 |  | 100 |  |
| Blot time [s] | 10 |  | 10 |  | 10 |  |
| **Microscopy** |  |  |  |  |  |  |
| Voltage | 300 |  | 300 |  | 300 |  |
| TEM mode | EFTEM<br>Nanoprobe |  | EFTEM<br>Nanoprobe |  | EFTEM<br>Nanoprobe |  |
| Indicated magnification | 105,000 |  | 105,000 |  | 105,000 |  |
| C2 aperture [μm] | 50 | | 50 | | 50 | |
| Spot size | 4 |  | 4 |  | 4 |  |
| Beam diameter [μm] | 1.55 | | 1.41 | | 1.45 | |
| Grid type | Quantifoil<br>R1.2/1.3<br>Cu/Rh 200 |  | Quantifoil<br>R1.2/1.3<br>Cu/Rh 200 |  | UltrAuFoil<br>R1.2/1.3 Au<br>300 |  |
| Objective lens aperture/phase palte [μm] | VPP | none | 100 | 100 | 100 | none |
| Target defocus [μm] | 0.4 | 0.7 – 1.2 | 0.8 – 1.5 | 0.8 – 1.5 | 0.6 – 1.4 | 0.6 – 1.4 |
| Zero-loss slit width [eV] | 25 | 25 | 25 | none | 25 | 25 |
| Pixel size [$\text{\AA}$] | 0.83 | | 0.83 | | 0.83 | |
| Super resolution | No |  | No |  | No |  |
| Exposure rate [e/pix/s] | 11.75 |  | 15.3 |  | 15 |  |
| Exposure rate [$\text{e}/\text{\AA}^2/\text{s}$] | 17.2 | | 22.2 | | 21.8 | |
| Exposure time [s] | 3.72 |  | 3.0 |  | 3.0 |  |
| Total exposure [$\text{e}/\text{\AA}^2$] | 64.0 | | 66.5 | | 65.4 | |
| Movie frames | 62 |  | 75 |  | 75 |  |
| Exposure per frame [$\text{e}/\text{\AA}^2/\text{frame}$] | 1.03 | | 0.89 | | 0.87 | |

The acquisition of the PAC1R dataset was started with the VPP (Figure 1B). After approximately 4,000 micrographs, the phase plate was retracted and the acquisition continued with the conventional defocus method (Figure 1C). The VPP provided a significant contrast improvement that simplified the visual identification of individual protein molecules in the micrographs (Figure 1B). The GLP-1R-TAS datasets consisted of an initial subset acquired with zero-loss energy filtering (ZLF) (Figure 1E) followed by a subset without energy filtering (Figure 1F). Energy filtering noticeably improved the contrast (Figures 1E and F). The GLP-1R-GLP-1 dataset used a 100 μm objective lens aperture (OLA) for the first half of the micrographs (Figure 1H) and no aperture for the rest (Figure 1I). The aperture had no discernable effect on the contrast of the micrograph but it blocked high-angle crystal reflections by the gold support film that appeared as bright spots in the images without an aperture (Figure 1I, arrow). In addition to parameters that were modified during the experiment, we also investigated the effects of defocus magnitude (DEF) and total exposure (EXP) by dividing the GLP-1R-GLP-1 dataset into corresponding subsets.

The VPP, ZLF and EXP subsets were processed independently through the complete single particle workflow in Relion (Zivanov et al., 2018). The DEF and OLA subsets were separated after the CTF refinement step in the processing of the complete GLP-1R-GLP-1 dataset. For each subset, we calculated several quantitative parameters (Table 2). Global 3D map resolution remains the most popular performance indicator in cryo-EM structural studies, mainly because of its convenience. However, it provides virtually no information about the behavior of the data. The B-factor is a more comprehensive measure of the overall performance. It models the combined effect of performance-reducing factors related to the sample, the experiment, and the data processing, as a Gaussian dampening function in reciprocal space (Rosenthal and Henderson, 2003). Higher values indicate stronger dampening and therefore lower performance. To determine the B-factor, the squared reciprocal resolution of independent reconstructions from random particle subsets of varying size is plotted as a function of the logarithm of the number of particles in the subsets. The B-factor is then equal to twice the reciprocal slope of the linear fit through the data points. We measured the B-factors of all subsets before and after particle polishing in Relion (Table 2, Figure S1). In many cases, the slope of the B-factor line, and therefore the B-factor did not change, although there was an improvement in resolution. To quantify the impact of such parallel shifts, we calculated two additional performance measures - the resolution from a fixed size 100k particle subset and the number of particles necessary to reach 3 Å resolution (Table 2). The two criteria are mathematically related to each other through the B-factor. Nevertheless, they provide two different viewpoints on the performance and therefore we included both in the analyses.

**Table 2.** Data processing details. Unless the row title indicates otherwise, values in brackets are the estimated standard error of the value.

|  | *PAC1R* |  | *GLP-1R-TAS* |  | *GLP-1R-GLP-1* |  | *GLP-1R-GLP-1* |  | *GLP-1R-GLP-1* |  |
| --- | --- | --- | --- | --- | --- | --- | --- | --- | --- | --- |
| | +VPP | -VPP | +ZLF | -ZLF | +OLA | -OLA | Def.<br>> 1 $\mu\text{m}$ | Def.<br>< 1 $\mu\text{m}$ | Exp.<br>65 e/$\text{\AA}^2$ | Exp.<br>40 e/$\text{\AA}^2$ |
| **Data processing** |  |  |  |  |  |  |  |  |  |  |
| Micrographs | 4,032 | 3,617 | 5,508 | 3,251 | 3,003 | 2,736 | 2,772 | 2,967 | 5,739 | 5,739 |
| Micrographs after CTF<br>fits (retention) | 3,761<br>(93%) | 3,553<br>(98%) | 4,839<br>(88%) | 2,638<br>(81%) | 2,177<br>(72%) | 2,490<br>(91%) | 2,120<br>(76%) | 2,547<br>(86%) | 4,667<br>(81%) | 4,596<br>(80%) |
| Measured defocus [$\mu\text{m}$] | 0.4 – 1.2 | 0.8 – 1.7 | 0.7 – 1.8 | 0.7 – 1.7 | 0.6 – 1.4 | 0.6 – 1.4 | 1.0 – 1.4 | 0.6 – 1.0 | 0.6 – 1.4 | 0.6 – 1.4 |
| Picked particles [$\times 10^3$]<br>(per micrograph) | 2,446<br>(650) | 3,012<br>(848) | 3,246<br>(671) | 1,490<br>(565) | 1,740<br>(799) | 2,046<br>(822) | 1,799<br>(849) | 1,987<br>(780) | 3,786<br>(811) | 2,875<br>(626) |
| Particles after<br>classification [$\times 10^3$]<br>(retention) (per<br>micrograph) | 553<br>(23%)<br>(147) | 607<br>(20%)<br>(171) | 181<br>(5.6%)<br>(37) | 140<br>(9.4%)<br>(53) | 297<br>(17%)<br>(136) | 338<br>(17%)<br>(136) | 318<br>(18%)<br>(150) | 318<br>(16%)<br>(125) | 636<br>(17%)<br>(136) | 618<br>(21%)<br>(134) |
| Resolution after CTF<br>refinement [$\text{\AA}$] | 3.12 | 2.83 | 2.83 | 2.99 | 2.35 | 2.33 | 2.37 | 2.31 | 2.23 | 2.44 |
| B-factor after CTF<br>refinement [$\text{\AA}^2$] | 137.5 (6.2) | 120.1 (5.3) | 124.0 (2.4) | 124.3 (6.9) | 72.9 (1.4) | 72.8 (1.6) | 75.5 (1.4) | 71.2 (2.3) | 73.6 (1.3) | 86.8 (2.0) |
| Particles to reach 3 $\text{\AA}$<br>after CTF refinement<br>[$\times 10^3$] | 958.0 (532) | 241.5 (127) | 78.0 (15) | 150.1 (91) | 22.5 (4.7) | 24.4 (5.7) | 25.8 (5.3) | 21.8 (7.6) | 22.9 (4.6) | 45.2 (12.0) |
| Resolution from 100k<br>particles after CTF<br>refinement [$\text{\AA}$] | 3.58 (0.18) | 3.22 (0.14) | 2.95 (0.04) | 3.09 (0.14) | 2.56 (0.05) | 2.58 (0.06) | 2.61 (0.05) | 2.55 (0.08) | 2.57 (0.05) | 2.78 (0.06) |
| **Resolution after**<br>**polishing [$\text{\AA}$]** | **2.99** | **2.69** | **2.72** | **2.87** | **2.27** | **2.25** | **2.25** | **2.23** | **2.14** | **2.2** |
| **B-factor after polishing**<br>**[$\text{\AA}^2$]** | **146.5 (3.4)** | **121.3 (2.8)** | **122.4 (4.9)** | **123.0 (3.5)** | **72.2 (1.1)** | **71.8 (1.6)** | **73.8 (1.7)** | **71.7 (1.5)** | **74.6 (1.2)** | **73.6 (0.8)** |
| Particles to reach 3 $\text{\AA}$<br>after polishing [$\times 10^3$] | 487.0 (132) | 116.7 (30) | 41.1 (17) | 71.9 (21) | 13.3 (2.2) | 14.0 (3.3) | 15.4 (3.8) | 12.6 (2.8) | 11.7 (2.1) | 17.7 (2.1) |
| Resolution from 100k<br>particles after polishing<br>[$\text{\AA}$] | 3.34 (0.07) | 3.04 (0.06) | 2.82 (0.07) | 2.93 (0.06) | 2.45 (0.03) | 2.45 (0.05) | 2.49 (0.05) | 2.43 (0.04) | 2.43 (0.04) | 2.51 (0.03) |

Besides improved contrast, the VPP did not provide any performance benefits. Conversely, the VPP maps had ~0.3 Å lower resolution than the conventional defocus maps (Figures 2A, S2A and S2C). The B-factor with the VPP was also significantly higher, by ~20 % (Figures 2B, S1A and S2B). The VPP required more than four times the number of particles to reach 3 Å resolution (Figures 2C and S2D). The VPP had the strongest impact on performance among all tested parameters. Unfortunately, it was detrimental by all indicators. This observation corroborates recent measurements of the attenuation of high-resolution signals by the VPP (Buijsse et al., 2020). The signal loss is stronger than the loss of electrons due to scattering, which is observable in Figure S3A, but the exact cause of the additional loss remains unknown.

**Figure 2.**
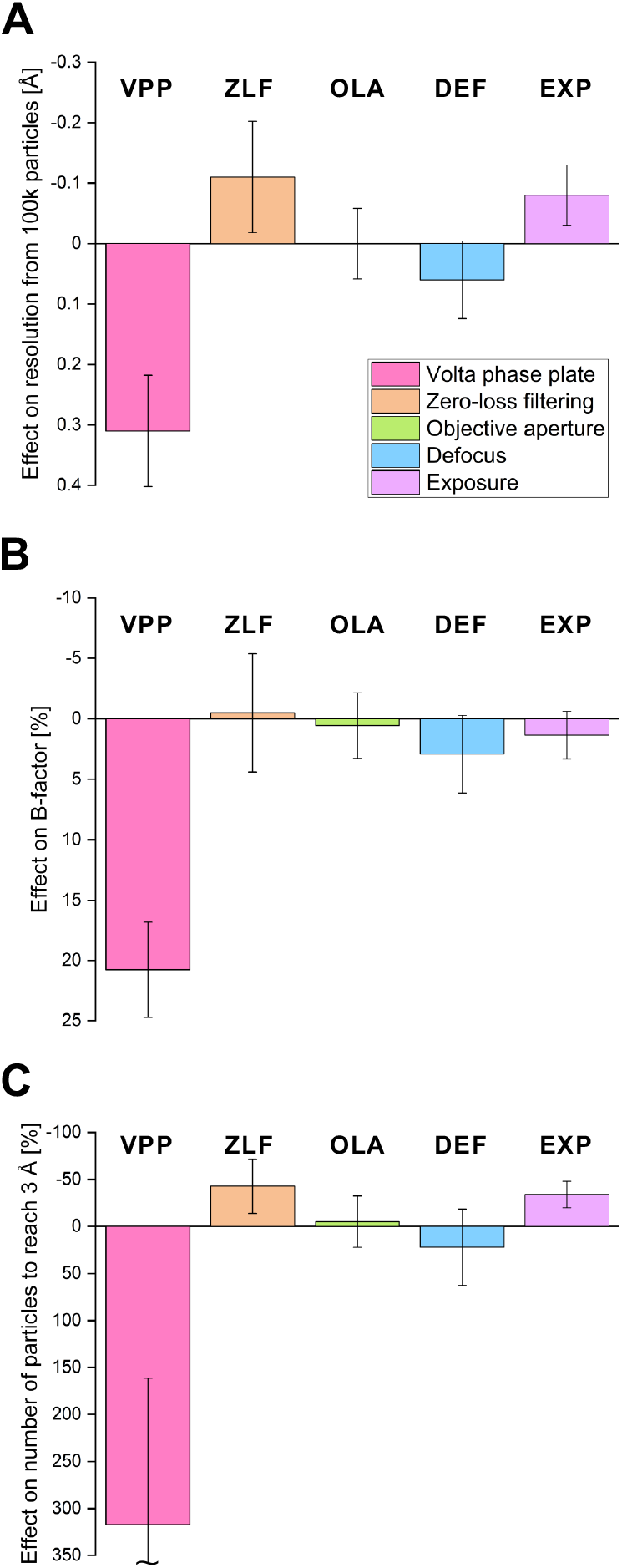
Effect of each experimental parameter on the three main performance measures. In all graphs, an upward bar indicates an improvement in performance. (A) Effect of the experimental parameters on the resolution from 100k particles expressed as the difference in Å. (B) Effect on the B-factor expressed as the change in % from the B-factor without the device, low DEF or low EXP to the B-factor with the device, high DEF or high EXP. (C) Effect on the number of particles to reach 3 Å resolution expressed as the change in % from the number of particles without the device, low DEF or low EXP to the number of particles with the device, high DEF or high EXP. Error bars represent the standard error of each value estimated from the statistics of the B-factor linear fits.

ZLF had an overall positive effect on the performance. It improved the resolution by ~0.1 Å (Figures 2A, S1B and S2A and S2C) and reduced the number of particles required to reach 3 Å approximately in half (Figures 2C and S2D). Surprisingly, ZLF had a negligible effect on the B-factor (Figures 2B, S1B and S2B). This would indicate that it improved the signal-to-noise-ratio (SNR) uniformly across all spatial frequencies. Indeed, one of the primary effects of ZLF is removal of inelastically scattered electrons that do not contribute to the image contrast and add background noise (Sigworth, 2016). However, in this case the sample thickness was only ~17.5 nm (Figure S3D) and the difference between the average image intensity with/without ZLF is ~1.7 counts/pixel (~5 %) (Figure S3C). Therefore, noise reduction alone cannot explain the observed performance gain. Most likely, it is due to additional amplitude contrast signal generated by ZLF (Yonekura et al., 2006). Such contribution is also supported by the perceptibly better contrast in ZLF images (Figures 1E versus 1F). Our results recapitulate the improvement observed in a recent sub-2 Å cryo-EM reconstruction of a homopentameric GABA_A_ receptor (Nakane et al., 2020). However, unlike the results in that study, we did not detect an improvement in the B-factor, possibly due to the wider energy limiting slit of 25 eV in our experiments compared to 5 eV in the GABA_A_R case.

The objective lens aperture had no impact on the performance. Differences in the quantifier values are minor and can be attributed to statistical fluctuations. The resolution values after polishing were identical, the B-factors were within 0.6 % and the number of particles necessary to reach 3 Å were within 4 % (Table 2, Figures 2, S1C and S2). Interestingly, in all but the 100k particles resolution test (Figure S2C) the subset without an aperture showed slightly better results. The effect of the aperture was clearly visualized in the electron count histograms of the subsets (Figure S3E) where it reduced the average intensity by ~0.5 counts/pixel (~1.4 %). This confirms that it did intercept high-angle scattered electrons, but the fraction was too small to have a detectable effect on contrast and performance.

Defocus is the main phase contrast mechanism in cryo-EM. It is a deliberately introduced aberration that delocalizes and phase shifts signals proportionally to their spatial frequency (Sigworth, 2016). When such signals are mixed with the primary unscattered wave, they create intensity modulation in the image that can be detected by the camera. Higher defocus values cause broader delocalization and visualize lower frequencies, thereby increasing the overall contrast. During the experiment setup, there is a natural urge to use higher defocus because it makes the molecules of the sample easier to see in the images. However, during processing, strongly delocalized high-resolution signals require more care and are more difficult to recover (Tegunov et al., 2020). Our experience showed that using lower defocus values, and the associated much lower image contrast, did not lessen the ability to locate and accurately align the particles during reconstruction. On the contrary, reconstructions from datasets collected with lower defocus, where particles were barely distinguishable in the images, produced higher resolution reconstructions. To test the impact of different defocus amounts, we split the GLP-1R-GLP-1 dataset into two equal halves, after the CTF refinement step, and independently processed them further. The split point happened to be very close to 1 μm defocus (1.023 μm). The low defocus subset produced slightly higher resolution reconstruction (2.23 vs 2.25 Å), had a lower B-factor (71.7 vs 73.8 Å2) and required 22% less particles to reach 3 Å (Figures 2, S1D and S2). The most important outcome from this test was that the low defocus subset did not show any performance deficits, despite the very low contrast in the images. This result suggests that with optimized samples in thin ice, the low frequency components are not essential for achieving accurate translational and rotational alignments. Furthermore, the slight advantage of low defocus persisted for the small particle number/lower resolution datapoints in the B-factor plot (Figure S1D, 4k-16k particles) where any high-resolution impairment of the high defocus data due to optical effects or inadequate processing should not play a role. We used a particle box size (320 pixels/265.6 Å) that was large enough to accommodate delocalized signals from the center of the particle at the final resolution (2.25 Å) and the highest measured defocus (1.4 μm). In regions away from the center of the box, however, there may be loss of high-resolution information due to delocalization. To test if this was the case, we re-extracted the particles without polishing with a 400 pixels/332 Å box, which should accommodate delocalized signals from all well resolved portions of the complex. The B-factor plot indeed showed a slight improvement of the high defocus relative to the low defocus line (Figure S2D). However, the B-factor ratio remained the same and the absolute resolutions for both subsets decreased due to the overall reduced signal-to-background ratio in the larger box. We did not deem it necessary to perform the resource-hungry polishing of the larger box particles because this is unlikely to provide any valuable information.

The last experimental parameter that was tested was the total exposure. There is no consensus in the community about the exposure threshold past which there will be no further gain in the quality of reconstructions. It will depend on the amount of signal in each projection, hence the size of the particle, with smaller molecules possibly benefiting more from higher exposures (Grant and Grigorieff, 2015). The direct detector movie acquisition format combined with exposure weighting or Bayesian polishing provides optimal signal extraction from the frame stack (Zivanov et al., 2019). In principle, this means that images can be overexposed indefinitely. However, longer exposures reduce the acquisition, data storage and data processing throughputs and one must make a practical compromise. Nowadays, the exposure used for single particle analysis is typically in the range of 30 to 80 e/Å^2^. To test the effect of total exposure on the performance for GPCRs, we re-processed the GLP-1R-GLP-1 dataset, from the beginning, using a 40 e/Å^2^ (46 frames) subset from the original 65 e/Å^2^ (75 frames) movies. The results showed a slight decrease in resolution and B-factor with the lower exposure (Table 2, Figures 2, S1E and S2). The 40 e/Å^2^ dataset produced a 0.08 Å lower resolution from 100K particles and needed 34 % more particles to reach 3 Å (Table 2, Figure 2). This indicates that there is a noticeable benefit from using higher total exposures (>60 e/Å^2^) for GPCRs. It is not clear why the increase from 40 to 65 e/Å^2^ had a positive effect at high resolutions. In the wake of radiation damage, only low-resolution features of the sample (>7 Å) would be contributing to the frames beyond 40 e/Å^2^ (Grant and Grigorieff, 2015). For the same reason, we expected that low defocus particles, that already have weak overall contrast, will be affected more by limiting the exposure. Curiously, this was not the case and lower defocus particles were a slightly bigger fraction of the final particle set in the 40 e/Å^2^ dataset (Figure S1F). Also unexpected was the significantly higher B-factor (86.8 vs 73.6 Å^2^) of the 40 e/Å^2^ dataset before polishing (Figure S1E, dashed lines). Motion correction statistics (Figure S5C) showed higher early motion and total motion with the lower exposure, indicating that the initial motion correction was probably less accurate. Despite this, particle polishing later in the processing was able to correct the discrepancy, as shown by the improved final result (Figure S1E). Better effectiveness of the initial multi-patch motion correction at high total exposures could also explain the unchanging B-factor after polishing for most of the subsets (Table 2, Figure S1). Interestingly, the VPP exhibited an opposite effect. The B-factor increased noticeably after polishing (146.5 vs 137.5 Å^2^) due to bigger signal gains at low spatial frequencies (Table 2, Figure S1A). This would suggest a bias of the motion tracking during polishing towards the much stronger low frequency components visualized by the phase plate.

Unsurprisingly, the quality of frozen grids had a significant impact on the outcome of GPCR experiments. High purity, good homogeneity and sufficient concentration of the protein solution were essential for achieving good results. To this end, all GPCR protein preparations were characterized for purity, assembly state and stability by biochemical assays and negative stain electron microscopy (Liang et al., 2017, 2020). During the initial experiments, we screened the cryo-EM grid plunging conditions, such as glow discharge time, blot force and blot time for our combination of a plasma cleaner and a plunging device. After determining a set of parameters that consistently produced thin ice over large portions of the grid, we kept these parameters constant. We had to significantly extend the glow discharge time to 90 s (Table 1), to improve grid wettability and enhance sample drainage during blotting. This reduced the pooling of solution in the middle of grid squares, especially with 200 mesh grids (compare Figures S4A, S4B with S4C, S4D). We also settled on a relatively long blotting time of 10 s that produced uniformly thin ice more consistently (Table 1). For each new sample, we only screened the concentration by preparing 2-3 grids with 2x dilution in-between. In our experience, GPCR samples produce optimal grids at concentrations between 3 and 7 mg/ml. The highest resolution results came from grids with uniformly thin ice that contained a single layer of molecules covering 50 – 90 % of the image area (Figure 1, Figure S4).

Figure 3 shows the history of our GPCR observations in the past 17 months. There is a general trend towards better resolution with a pronounced jump in August 2019. At that point, we started using gold foil support grids (Russo and Passmore, 2016) instead of the holey carbon grids that were used in all previous experiments (Table 1). This improved the resolution substantially, on average by ~0.5 Å, which was very beneficial but also quite unexpected. Previously, we had achieved 1.62 Å resolution with an apoferritin test sample on the same microscope by using holey carbon grids (Danev et al., 2019). Therefore, we did not suspect that carbon film grids were imposing a performance penalty with the GPCR samples, where the resolution was typically in the 2.5 – 3.0 Å range (Figure 3). Nevertheless, the improvement from the gold foil grids was evident and was confirmed consistently by the followup experiments (Figure 3). Quantitative analysis of the ice thickness in the GLP-1R-TAS and GLP-1R-GLP-1 datasets showed that it was very similar and in the range of 150-200 Å (Figures S3D and F). Therefore, the resolution improvement was not due to thinner ice. The PAC1R dataset had a much wider thickness distribution that extended towards thicker ice (Figure S3B), which could explain its slightly lower resolution. Beam-induced motion statistics showed a clear reduction in the early motion with the Au foil grid (Figure S5A) but similar total motion (Figure S5B). The early stage of an exposure carries the highest resolution signal because it has the lowest radiation damage. This, and the expected vertical doming of the ice layer with carbon film supports (Brilot et al., 2012) could explain the strong benefit that we observed from using gold foil grids. Since switching to such grids, we are routinely getting resolutions at or below 2.5 Å in our reconstructions though this remains dependent upon sample quality (Figure 3).

**Figure 3.**
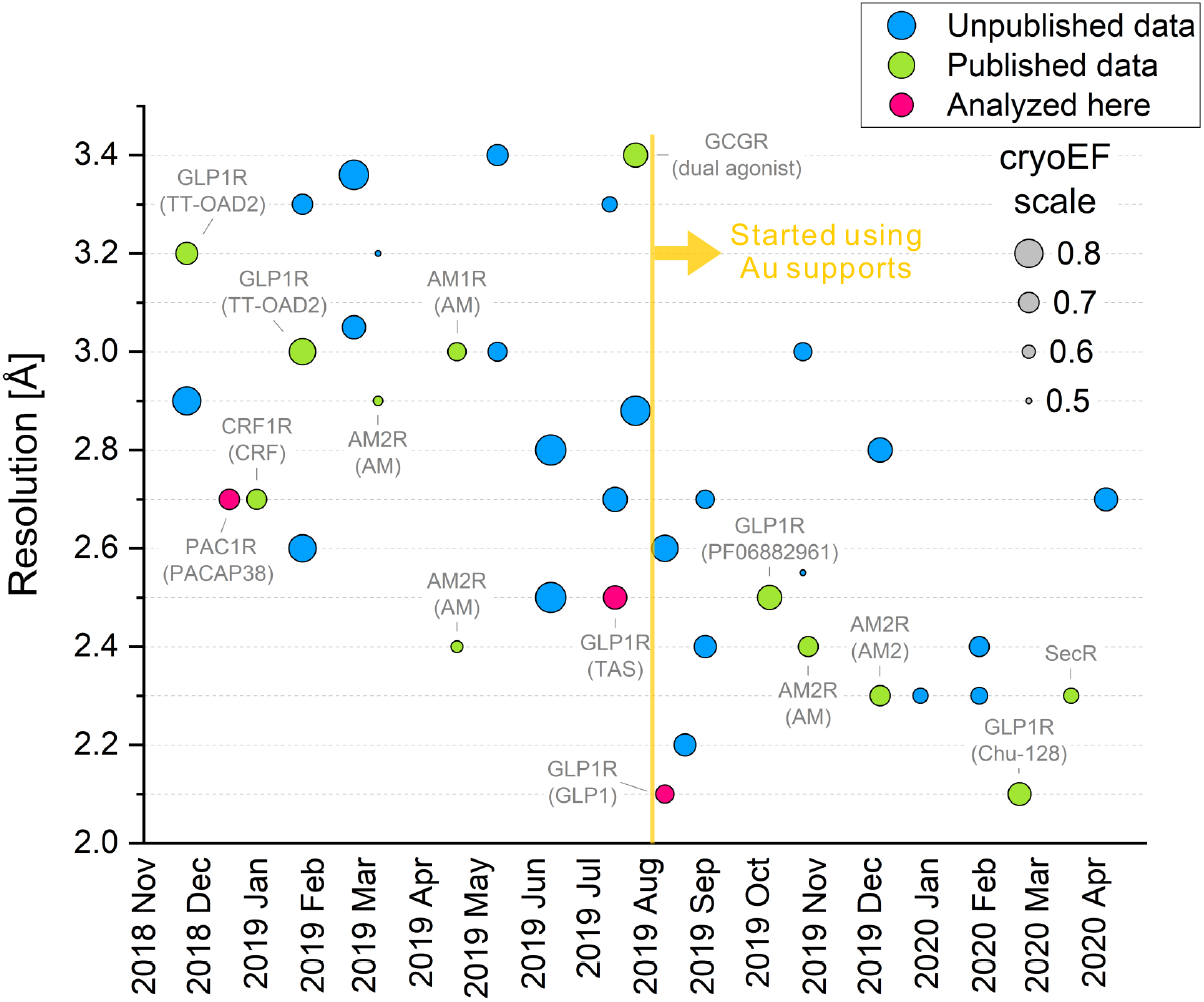
History of our cryo-EM GPCR reconstructions. Over the year and a half period shown in the plot, there is a general trend towards better resolution with a significant improvement in August 2019, when the sample supports were switched from holey carbon films to gold foil grids. The size of each dot represents the cryo-EM efficiency quantity (see legend) that estimates the uniformity of the particle orientation distribution. A value of 0.5 corresponds approximately to an angular distribution that is barely sufficient for producing a usable 3D map and a value of 1 corresponds to an ideal uniform distribution. Published results are annotated and the three datasets from this study are highlighted in pink.

To further characterize the behavior of samples, we calculated the cryo-EM “efficiency” (cryoEF) (Naydenova and Russo, 2017) for all datasets (Figure 3, dot size). CryoEF measures the uniformity of the particle orientation distribution and values above 0.5 are considered to indicate that a dataset will produce a usable 3D map. All datasets in Figure 3 satisfied this criterion, with only four of them having cryoEF values between 0.5 and 0.6 and the majority in the range 0.65 to 0.75. There was no strong correlation between cryoEF and map resolution. The transition to gold foil grids appears to have truncated high (>0.75) cryoEF values without a negative impact on resolution, possibly because of consistently thinner ice that excludes particles oriented with their long axis perpendicular to the specimen plane.

Figure 4 shows a similar region from the PAC1R and the GLP-1R-GLP-1 maps and models to illustrate the increase in map fidelity from 2.7 Å to 2.1 Å. The most significant improvement is in the definition of sidechain densities, which increases the probability of correct rotamer orientation assignment, measuring interaction distances and identifying water molecules (Figure 4B). Such information can be crucial in the accurate interpretation of the interactions between the receptor and the ligand, which could be a peptide or a small molecule.

**Figure 4.**
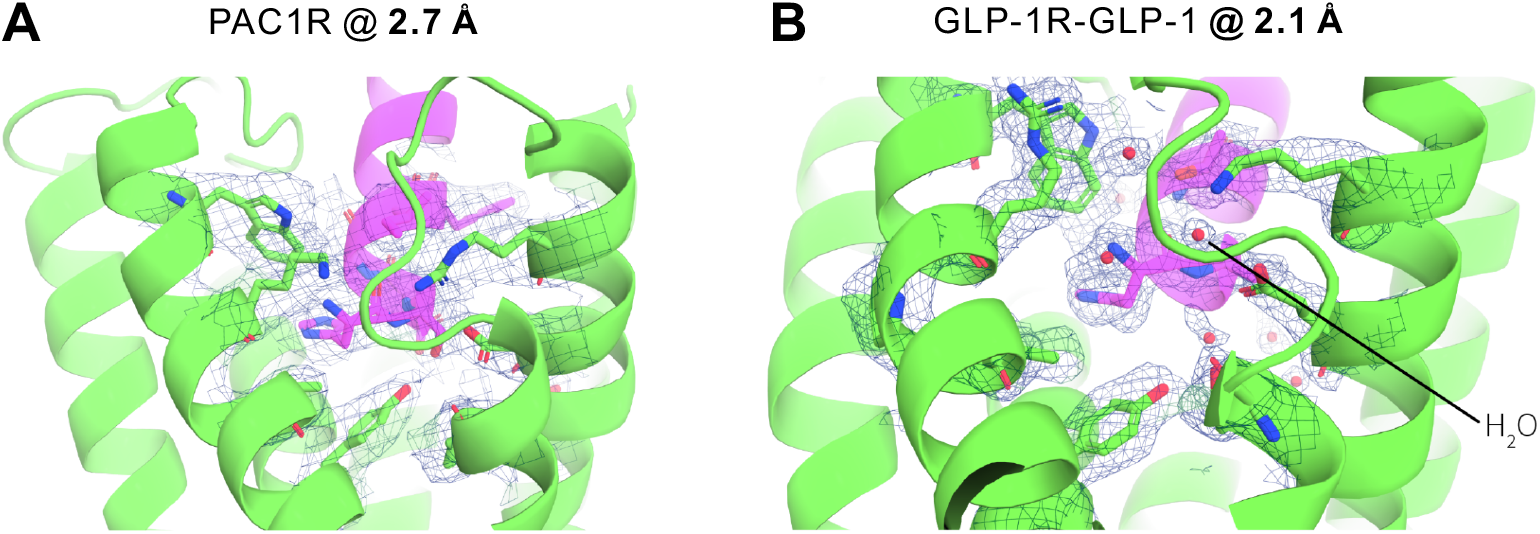
Illustration of the atomic modeling benefits from improved map resolution. Cartoon representation of the N-terminal portion of the agonist peptides GLP-1 and PACAP38 bound to their respective receptors. The cryo-EM density maps are drawn at 5-sigma as a blue mesh and clearly highlight the ability to more accurately model side chain and water positions at a higher spatial resolution. (A) N-terminus of the PACAP38 peptide (magenta) bound to the PAC1 receptor (green) at 2.7 Å resolution. There were no reliably detectable water molecule densities in this region of the map. (B) N-terminus of the GLP-1 peptide (magenta) bound to GLP-1 receptor (green) at 2.1 Å resolution. Several water molecule densities were identified and modelled inside the binding pocket of the receptor where they facilitate the interaction with the ligand.

## 3. Conclusions

Here, we presented a comprehensive investigation of the effect of several experimental parameters on the performance of cryo-EM single particle analysis for G protein-coupled receptors. The first major conclusion is that the Volta phase plate does not provide any benefits in such studies and therefore researchers should refrain from using it in their experiments. Except for sample optimization, the rest of the parameters had much smaller but nonetheless important contributions to the performance.

Zero-loss energy filtering improved the resolution by ~0.1 Å throughout the resolution range. The contribution appears to be due to additional amplitude contrast generated by removing inelastically scattered electrons. The improvement of high-resolution components may indicate that there could be other effects that will need further investigation.

The objective lens aperture had virtually no effect on the performance. With the gold foil grids it did prevent crystal reflection spots from appearing in images and therefore we consider it to be beneficial. Nonetheless, it should be used with care because it may cause wavefront distortion of high-resolution components (<2 Å) due to electrostatic potentials on the aperture.

Our results show that there is a practical advantage of using low defocus values (<1 μm) with optimized samples. Current particle handling strategies in the most popular image processing packages impose a restriction on the minimum particle box size as a function of the maximum defocus in the dataset that will preserve delocalized high-resolution components within the box. The recently introduced M package overcomes this limitation by performing a CTF sign correction on a larger area before extracting the particle box (Tegunov et al., 2020). This was shown to solve the resolution limitation of higher defocus values and could allow their use in the future without detrimental effects. We measured a ~0.06 Å resolution benefit from a using a higher 65 e/Å^2^ total exposure compared to 40 e/Å^2^. The difference was larger for non-polished particles and movie alignment statistics indicated that the merit from the higher exposure is most likely due to improved initial motion correction of the movies.

Sample optimization had a significant impact on the quality of results. High protein concentrations, to get a monolayer of molecules in thin ice, improved the quality and consistency of the samples. Gold foil grids had the largest positive impact among all tested parameters on the resolution of reconstructions; on average by ~0.5 Å. Their advantage appears to be a combination of reduced initial beam-induced motion and more consistent sample quality with uniformly thin ice.

Other factors, that were not tested in this study, will also have a substantial influence on the performance. Direct detector technology continues to evolve and recent innovations include an electron even representation (EER) format that mitigates the limitation of fixed movie frame width by recording every individual electron event (Guo et al., 2020). This and other developments will contribute to achieving better signal-to-noise ratio and higher throughout that will provide significant benefits in the future.

The experimental parameter optimization presented here can be applied to other similarly sized membrane proteins and will possibly result in comparable performance gains.

## 4. Methods

### Protein expression and purification

The expression and purification of the PAC1R (Liang et al., 2020), GLP-1R-TAS (in preparation) and the GLP-1R-GLP-1 (Zhang et al., 2020) was described in detail previously.

### Cryo-EM sample preparation

Sample support grids were washed in advance by placing them on a piece of filter paper in a glass Petri dish, soaking the paper with acetone and letting the solvent evaporate. Immediately before sample preparation, the grids were glow discharged in low pressure air with 10 mA current in a PIB-10 Ion Bombarder (JEOL, Japan). Cryo-EM grids were prepared by plunge-freezing in liquid ethane on a Vitrobot Mark IV (Thermo Fisher Scientific, USA). The glow discharge and plunge-freeze parameters are listed in Table 1.

### Cryo-EM data collection

The datasets were collected on a Titan Krios G3i (Thermo Fisher Scientifics, USA) 300 kV electron microscope equipped with a GIF Quantum energy filter and a K3 direct electron detector (Gatan, USA). Movies were acquired with homemade scripts in SerialEM (Schorb et al., 2019) using 9-hole beam-image shift acquisition pattern with 1 image in the center of each hole and saved as non-gain-normalized compressed TIFF files. The acquisition parameters for each dataset are listed in Table 1 and the dataset details in Table 2.

### Data processing

All datasets were motion corrected with MotionCor2 (Zheng et al., 2017) using 5 x 3 patches (long x short edge of the K3 image area), no frame grouping, B-factor 500, with saving of dose weighted and non-dose weighted averages. The CTFs were fitted on the non-dose-weighted averages using Gctf (Zhang, 2016) with 20 – 3.5 Å resolution range, 1024 pixel box, EPA averaging, high-resolution refinement with 15 – 3.0 Å resolution range, resolution cross-correlation cutoff limit of 0.5 and defocus search range determined from the defocus histogram of an initial trial CTF fit. For the PAC1R and GLP-1R-TAS datasets, a 20 A low-pass filtered GPCR map from a previous reconstruction was used for reference-based particle picking, a round of 2D classification and an initial round of 3D refinement in Relion 3 (Zivanov et al., 2018) the resulting 3D map from which was used as a reference and to create a 3D mask for the 3D classification rounds. For the GLP-1R-GLP-1 dataset, an initial round of reference-free picking, followed by 2D classification, initial-model generation by SGD (Punjani et al., 2017) and an initial round of 3D refinement in Relion 3, the resulting 3D map from which was used for reference-based particle picking of all micrographs, and as a reference and to make a mask for the 3D classification rounds. All three datasets were cleaned solely by 3D classification of the original set of picked particles. For the PAC1R VPP subset, two rounds of 3D classification with 3 and 2 classes were performed, each time selecting the best resolved class. For the PAC1R defocus subset, three rounds of 3D classification with 3, 3 and 2 classes were performed, each time selecting the best resolved class. For the GLP-1R-TAS zero-loss filtered subset, three rounds of 3D classification with 3, 3 and 3 classes were performed, each time selecting the best resolved class. For the GLP-1R-TAS non-energy-filtered subset, three rounds of 3D classification with 3, 3 and 3 classes were performed, each time selecting the best resolved class. For the GLP-1R-GLP-1 dataset, three rounds of 3D classification with 3, 3 and 5 classes were performed, each time selecting the best resolved class. The resulting particle sets after the 3D classification filtering steps are listed in Table 2. The +VPP, -VPP, +ZLF, and -ZLF subsets were each independently subjected to two rounds of per-image-shift-position CTF refinement, 3D auto refinement, particle polishing, and 3D auto refinement in Relion 3. The GLP-1R-GLP-1 complete dataset was subjected to two rounds of per-image-shift-position CTF refinement, 3D auto refinement, particle polishing, and 3D auto refinement in Relion 3. The +OLA, -OLA, high-, and low-defocus subsets were separated by re-extracting the particles from the micrographs with the CTF parameters from the last CTF refinement step, then 3D refined, polished independently and again 3D refined in Relion 3. The 40 e/Å^2^ exposure frame subset from the GLP-1R-GLP-1 dataset was processed independently through the complete workflow by using the first 46 of the 75 movie frames. The 40 e/Å^2^ frame subset was subjected to the same set of processing steps, including movie alignment, CTF fitting, and particle picking, to take into account the effect that reduced exposure may have in each of these steps. The result from the last 3D refinement of each subset was subjected to B-factor estimation with random particle subsets using the B-factor calculation Python script supplied with Relion 3. The refinement and B-factor results after the first CTF refinement step and after the last particle polishing step for each subset are presented in Table 2.

The standard errors of the values in Table 2 and Figures 2 and S2 were calculated from the standard errors of the slope and offset values of the B-factor linear fits in Figure S1.

The cryo-EM efficiency (cryoEF) value was calculated on the particle set from the final 3D refinement of each dataset using the cryoEF program (Naydenova and Russo, 2017) with 256 pixels box size.

The ice thickness in Figure S3 was calculated using inelastic mean-free path of 395 nm without an objective aperture and 322 nm with aperture (Rice et al., 2018).

## Acknowledgements

R.D. was supported by the Japan Society for the Promotion of Science (JSPS) KAKENHI #18H06043, Takeda Science Foundation 2019 Medical Research Grant and Japan Science and Technology Agency PRESTO (18069571). D.W. is a Senior Research Fellow of the Australian National Health and Medical Research Council (NHMRC). P.M.S. is a Senior Principal Research Fellow of the NHMRC. The work was supported by a Program grant (1150083) and Project grants (1120919, 1126857 and 1159006) from the NHMRC. M.J.B is supported by a US-DoD grant (PR180285, W81XWH-19-1-0126).

## Author contributions

R.D., D.W. and P.M.S. conceived the research. R.D. developed the method, collected and processed cryo-EM data, analyzed results and write the initial draft. M.B. processed cryo-EM data and analyzed results. Y.L.L. and X.Z. expressed and purified protein complexes, performed biochemical assays, negative stain screening and analyzed cryo-EM data. All authors participated in the editing of the manuscript.

---

**Figure S1.**
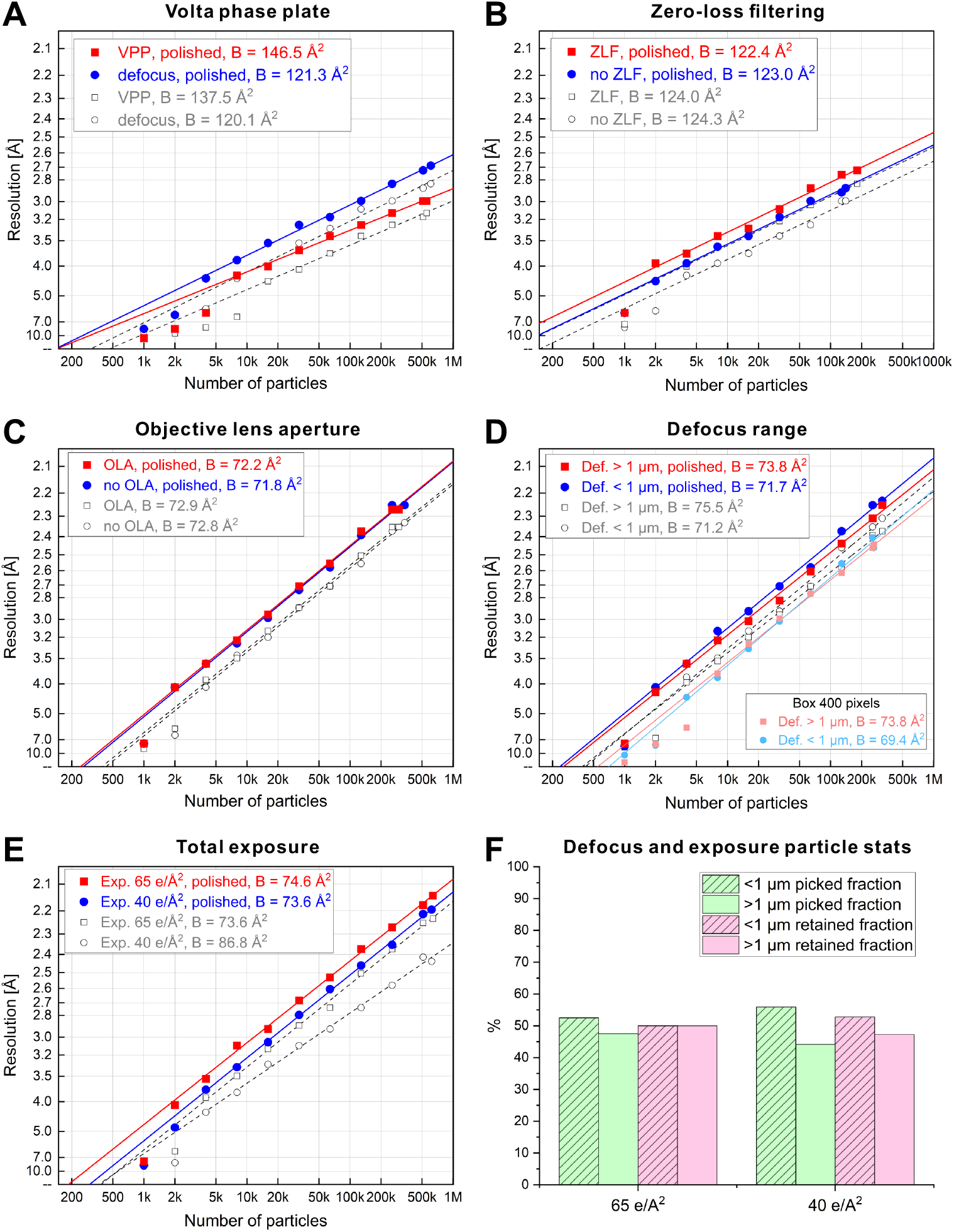
B-factor plots for all particle subsets and particle retention statistics for the total exposure subsets. (A) B-factor plots for the VPP/defocus subsets of the PAC1R dataset, before and after particle polishing. (B) B-factor plots for the zero-loss energy filtering and no-energy-filtering subsets of the GLP-1R-TAS dataset, before and after particle polishing. (C) B-factor plots for the objective lens aperture/no aperture subsets of the GLP-1R-GLP-1 dataset, before and after particle polishing. (D) B-factor plots for the below and above 1 μm defocus subsets of the GLP-1R-GLP-1 dataset, before and after particle polishing. Also included are B-factor plots for the defocus subsets reconstructed with a 400 pixels box, before particle polishing. (E) B-factor plots for the full and partial total exposure micrograph sets from the GLP-1R-GLP-1 dataset, before and after particle polishing. (F) Defocus range particle fractions as picked (green) and in the final particle set (pink) for the full and partial total exposure sets from the GLP-1R-GLP-1 dataset.

**Figure S2.**
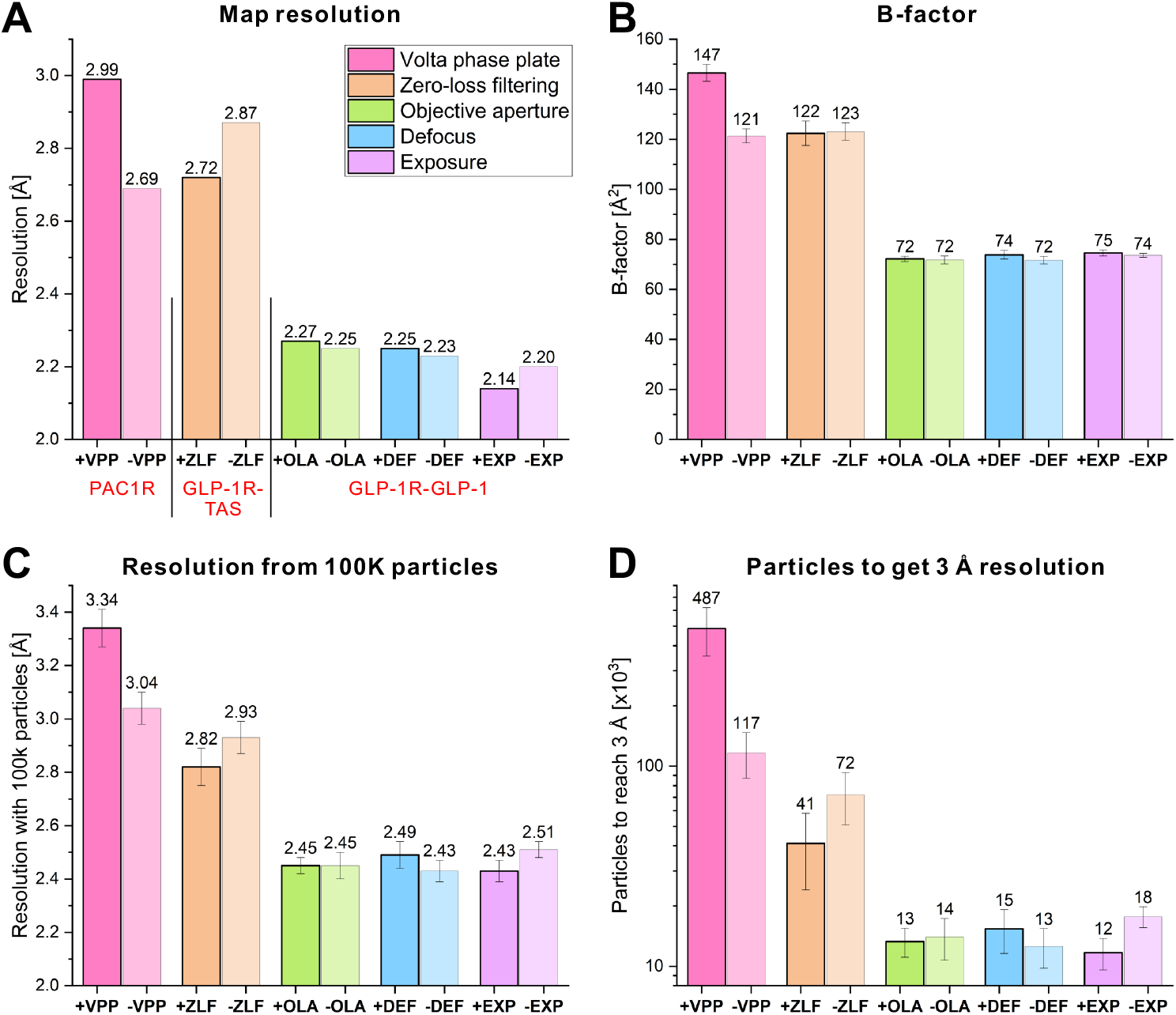
Graphical summary of the performance test results. (A) Resolution of the reconstructions of the complete particle subsets. The results are not directly comparable because the subsets have different particle numbers. (B) B-factor of each subset measured from the B-factor plots in Figure S1. The results for each sample are directly comparable. (C) Resolution from 100k particles in each subset. (D) Number of particles in each subset required to reach 3 Å resolution.

**Figure S3.**
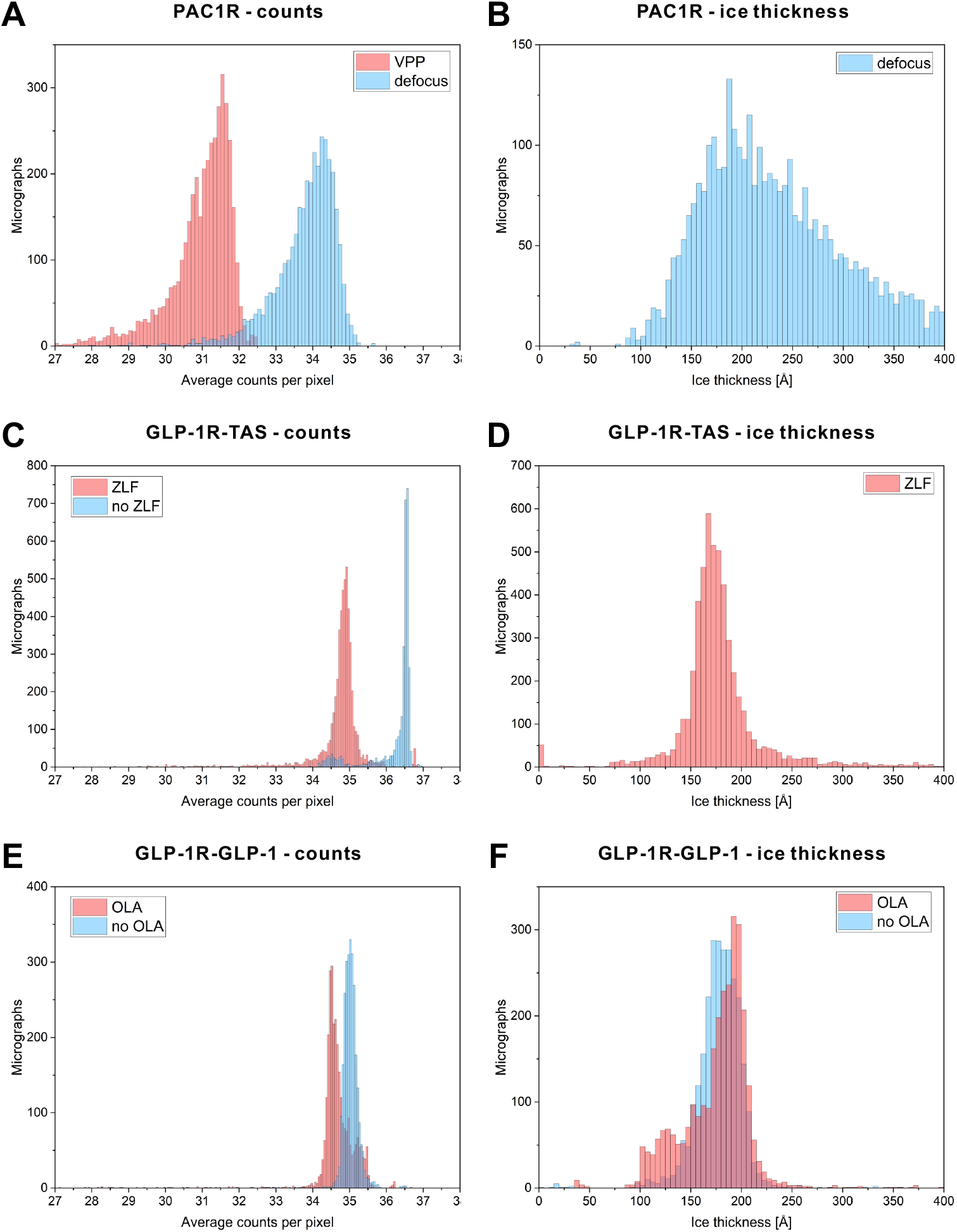
Electron event counts and ice thickness statistics for each dataset. (A) Average count distribution for all movies in the PAC1R dataset. The VPP reduced the average counts by ~10 % due to scattering. (B) Ice thickness distribution of the conventional defocus subset from the PAC1R dataset. The average ice thickness in ~200 Å but there is a long tail towards higher thickness, indicating a nonuniform or variable ice thickness distribution. (C) Average count distribution for the movies in the GLP-1R-TAS dataset. Zero-loss energy filtering reduced the intensity by ~5 % by removing inelastically scattered electrons. (D) Ice thickness distribution of the zero-loss subset of the GLP-1R-TAS dataset. The thickness is in the range of 150 – 200 Å with an average of ~175 Å. (E) Average count distribution of the GLP-1R-GLP-1 dataset. The objective lens aperture intercepted high-angle scattered electrons thus reducing the intensity by 1.4 %. (F) Ice thickness distribution of the GLP-1R-GLP-1 dataset. The thickness was very similar to that of the GLP-1R-TAS dataset in (D), with a range of 100 – 220 Å and an average of ~175 Å.

**Figure S4.**
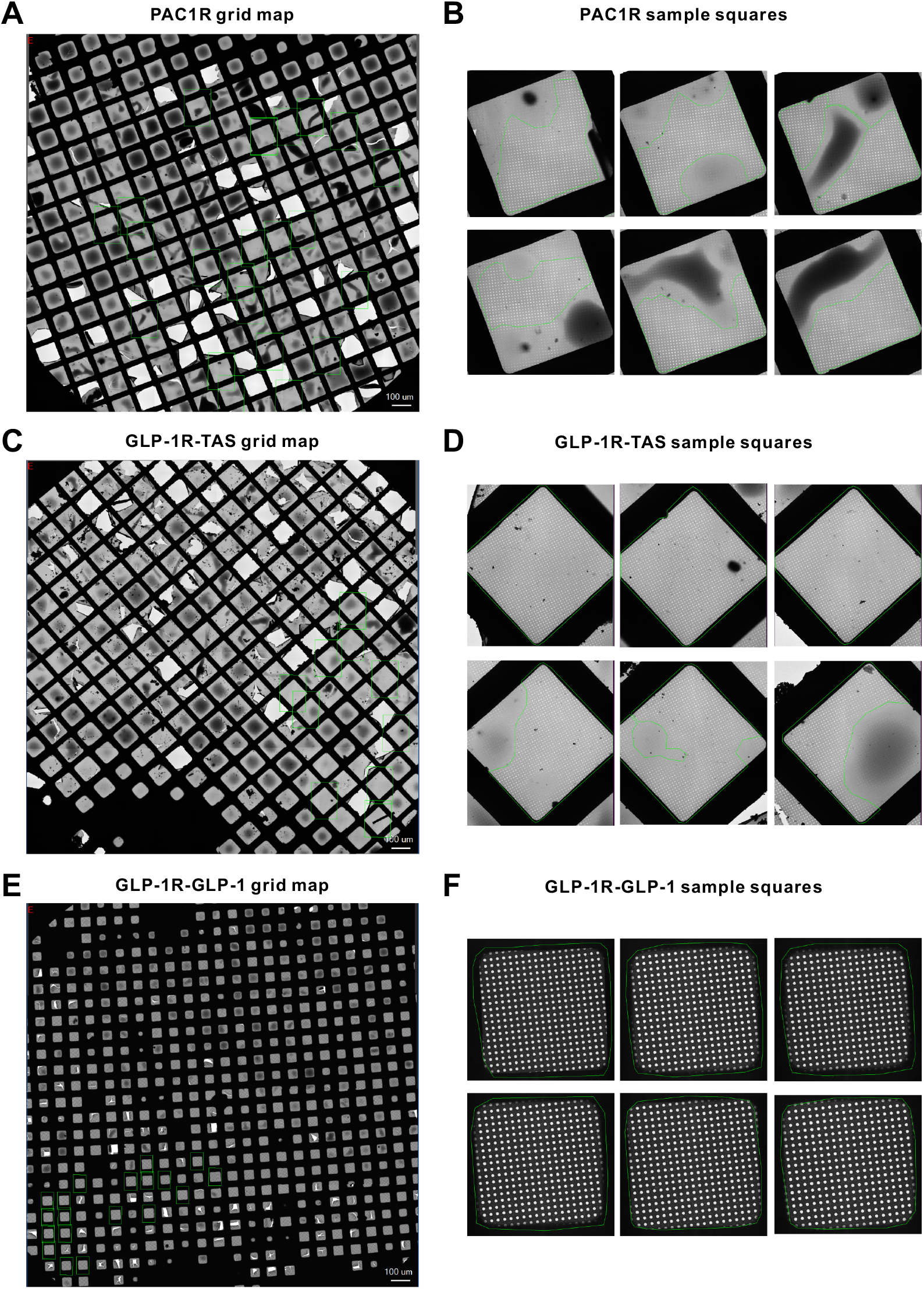
Grid maps and representative grids squares from the datasets. The PAC1R (A) and GLP-1R-TAS (C) datasets were collected using 200 mesh holey carbon grids. The GLP-1R-GLP-1 (E) datasets was collected using a holey gold foil 300 mesh grid. The 200 mesh carbon film grids (A and C) had more broken squares and more uneven ice distribution. The PAC1R (A) grid was glow discharged for 30 s, compared to 90 s for the GLP-1R grids (C and E). This reduced the drainage of the sample and caused more pooling of sample solution in the middle of squares (A and B versus C and D). The gold foil grid (E and F) had a more uniform ice thickness distribution and less broken squares. Green rectangles in (A, C and E) indicate the squares that were used for data acquisition. Green polygons in (B, D and F) encompass data acquisition areas on the shown squares.

**Figure S5.**
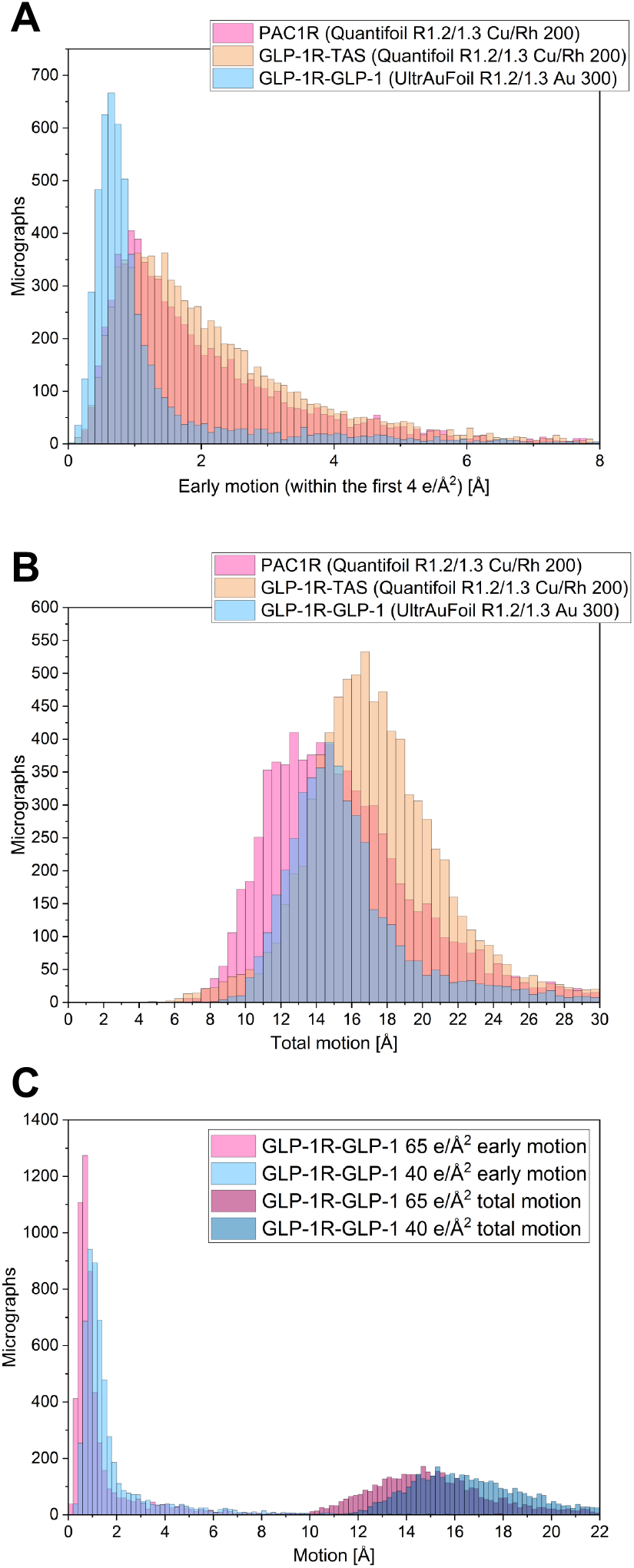
Motion statistics from the initial movie motion correction of the datasets. (A) Accumulated motion from the first 4 e/Å^2^ of the exposure. The GLP-1R-GLP-1 dataset (light blue) collected with a holey gold foil gird exhibited significantly less movement in the beginning of the exposure, which contributed to higher final map resolution (2.1 Å). The PAC1R (pink) and GLP-1R-TAS (beige) had broader initial movement distributions and produced maps at lower resolutions (2.7 and 2.5 Å). (B) The total accumulated motion had a similar distribution for the three datasets. (C) Motion statistics for the complete GLP-1R-GLP-1 65 e/Å^2^ exposure (pink) and an initial frame subset with 40 e/Å^2^ exposure (blue). The lower exposure subset had higher early and total motions that indicate a less effective motion correction with the lower total exposure.

